# A rich conformational palette underlies human Ca_V_2.1-channel availability

**DOI:** 10.1101/2024.09.27.615501

**Authors:** Kaiqian Wang, Michelle Nilsson, Marina Angelini, Riccardo Olcese, Fredrik Elinder, Antonios Pantazis

## Abstract

Depolarization-evoked opening of Ca_V_2.1 (P/Q-type) Ca^2+^-channels triggers neurotransmitter release, while voltage-dependent inactivation (VDI) limits channel availability to open, contributing to synaptic plasticity. The mechanism of Ca_V_2.1 response to voltage is unclear. Using voltage-clamp fluorometry and kinetic modeling, we optically tracked and physically characterized the structural dynamics of the four Ca_V_2.1 voltage-sensor domains (VSDs). VSD-I seems to directly drive opening and convert between two modes of function, associated with VDI. VSD-II is apparently voltage-insensitive. VSD-III and VSD-IV sense more negative voltages and undergo voltage-dependent conversion uncorrelated with VDI. Auxiliary β-subunits regulate VSD-I-to-pore coupling and VSD conversion kinetics. Ca_V_2.1 VSDs are differentially sensitive to voltage changes brief and long-lived. Specifically the voltage-dependent conformational changes of VSD-I are linked to synaptic release and plasticity.

## Introduction

The Ca_V_2.1, or P/Q-type, voltage-gated Ca^2+^ channel, is the predominant Ca_V_ subtype in the brain and it plays a crucial role in synaptic transmission^1, 2, 3, 4, 5, 6, 7^. Presynaptic Ca_V_2.1 channels convert an electrical signal (action potentials) into a biochemical signal (Ca^2+^ entry), triggering neurotransmitter release (fig.1a). A prolonged depolarization or train of action potentials cause Ca_V_2.1 voltage-dependent inactivation (VDI). During VDI, channels enter a non-conductive state and are not available to mediate Ca^2+^ influx. This contributes to short-term depression, a form of synaptic plasticity that affects informational encoding^8, 9, 10, 11^ (fig.1b). Postsynaptic Ca_V_2.1 channels generate depolarization-induced local Ca^2+^ transients and are implicated in long-term depression, which underlies cerebellar learning^4, 12^. Mutational studies in mice suggest a role of Ca_V_2.1 in synaptic plasticity, spatial learning and memory; while variants of *CACNA1A*, the gene encoding the Ca_V_2.1 pore-forming subunit α_1A_, are associated with serious neurological disease^13, 14, 15^.

**Fig. 1.**
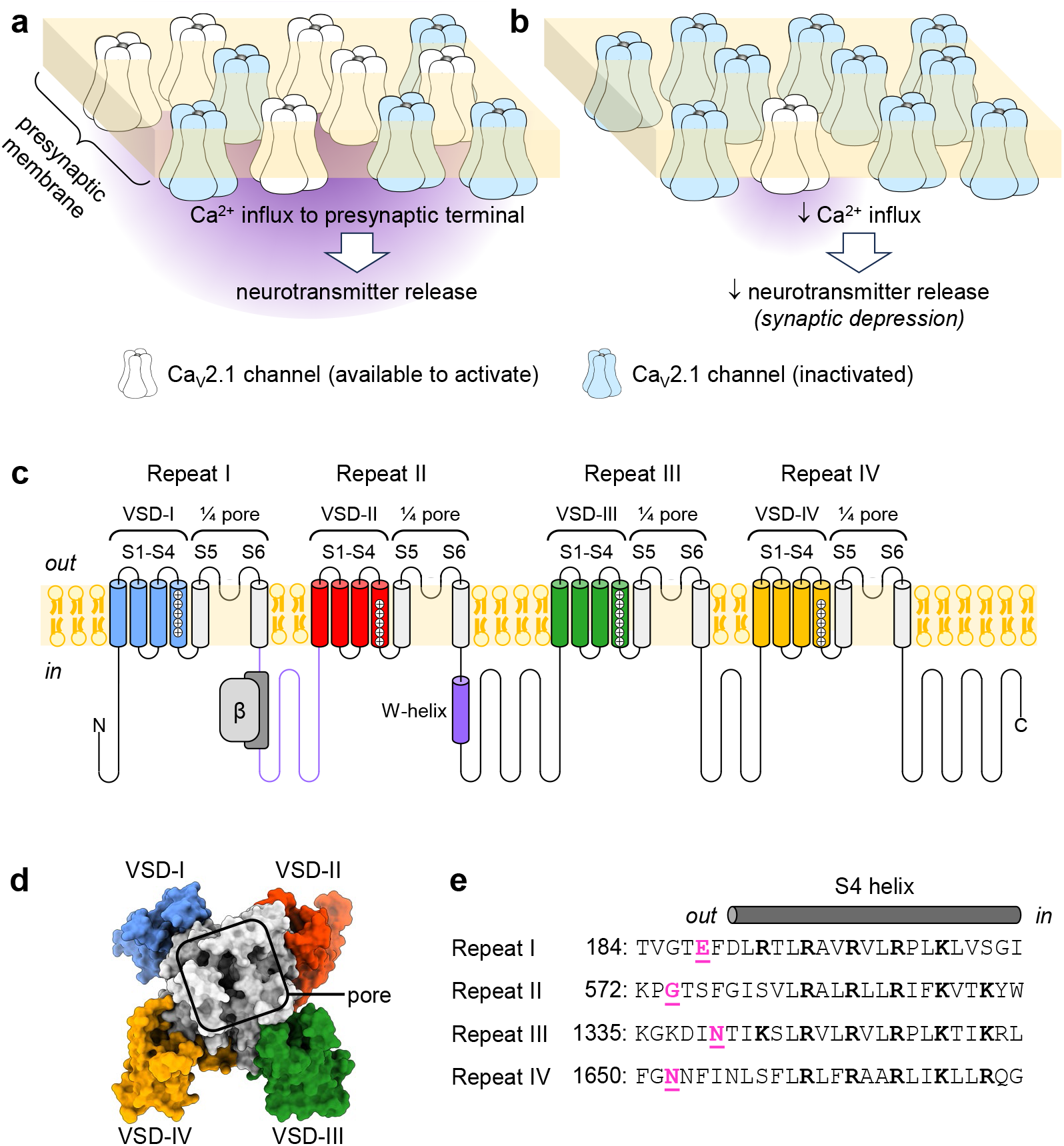
Ca_V_2.1 function and consequences of inactivation, its pore-forming subunit and its four non-identical VSDs. **(a)** Under normal conditions, some presynaptic Ca_V_2.1 channels are available to activate (white) in response to an action potential, mediating Ca^2+^ influx into the presynaptic terminus that triggers transmitter release. Some Ca_V_2.1 channels are inactivated (blue). **(b)** Prolonged depolarization or trains of action potentials induce voltage-dependent inactivation (VDI), further decreasing the number of available Ca_V_2.1 channels, and subsequently transmitter release. This contributes to synaptic plasticity. **(c)** The Ca_V_2.1 pore-forming subunit (α_1A_) contains four homologous repeats (I-IV). Membrane-spanning helices S1-S4 from each repeat comprise a voltage-sensor domain (VSD). The S5-S6 helices from each repeat form the ion-conducting pore. The auxiliary β-subunit binds between repeats I and II^45^. The intracellular I-II linker and W-helix within the II-III linker (indigo) act as blocking particles to occlude ion conductance during VDI^36, 37, 38, 39^. **(d)** Top view of α_1A_ (PDB: 8X90^31^). **(e)** S4 helix sequence comparison. Positively charged residues (bold) confer voltage sensitivity to the VSDs^16^. Amino-acid residues substituted to cysteine for fluorescence labelling in fig.2 are in magenta: VSD-I: E188; VSD-II: G574; VSD-III: N1340; VSD-IV: N1652.

Ca_V_2.1 channels consist of the α_1A_-subunit, extracellular α_2_δ and intracellular β-subunits^6, 7^. Channel voltage regulation stems from voltage-dependent conformational changes in the α_1A_ voltage-sensing apparatus^16, 17^. This comprises four transmembrane, homologous but non-identical, voltage-sensor domains (VSDs; fig.1c-e) but their roles in voltage-dependent activation and inactivation were not previously studied. Because Ca_V_2.1 has four different VSDs, it is possible that each VSD serves different functions to drive neurosecretion and contribute to synaptic plasticity. We optically tracked the voltage-dependent movements of the individual Ca_V_2.1 VSDs in conducting channel complexes in cellula by combining the cut-open oocyte vaseline gap voltage clamp^18, 19, 20^ with voltage-clamp fluorometry (VCF)^20, 21, 22, 23, 24^.

## Results

### Ca_*V*_*2*.1 VSDs activate with distinct voltage dependencies

To optically track the movements of individual VSDs under physiologically relevant conditions, we used VCF. Briefly, specific amino-acid residues at the extracellular loop between the S3 and S4 transmembrane helices of each repeat were mutated to cysteine (fig.1e). During the experiments, the engineered cysteine was modified with the thiol-reactive and environment-sensitive fluorophore, MTS-TAMRA. Thus the conformational rearrangements of the labelled VSDs in response to brief depolarizations were reported as ensemble fluorescence deflections (Δ*F*). To limit additional regulation (Ca^2+^ regulation or VDI), we (i) used Ba^2+^ as charge carrier, and pre-injected cells with the BAPTA Ca^2+^-chelator^17^; and (ii) studied Ca_V_2.1 channels including β_2a_, which slows down VDI relative to β-less channels^25, 26^.

VSD-I activated with a two-part voltage dependence (fig.2a,f): one component, “F1”, had a voltage-dependence very close to that of pore opening (calculated by normalized tail current, *I*_tail_), and the other (“F2”) was observed at very negative potentials.

We only detected faint Δ*F* signals (<0.1%) from VSD-II (fig.2b), similar in amplitude to Δ*F* from Ca_V_2.1 without a substituted cysteine (fig.2e), likely due to non-specific labelling. Lack of Δ*F* suggested that VSD-II does not undergo voltage-dependent conformational changes. In fact, no Δ*F* were detected despite (i) probing most of the S3-S4 linker, (ii) trying different fluorophores, (iii) removing a tryptophan that might quench the nearby fluorophore^27, 28, 29, 30^; (iv) using a different complement of auxiliary subunits; (v) neutralizing counter-charges that could stabilize the S4_II_ resting state^16^ and (vi) perturbing a PIP_2_ binding site, resolved in a Ca_V_2.1 structure^31^ (fig.S1). Indeed, none of the mutations used in the above studies significantly altered the voltage-dependence of pore opening (fig.S2), suggesting that VSD-II does not contribute to Ca_V_2.1 voltage sensitivity.

VSD-III and VSD-IV appeared to activate at negative potentials, close to the physiological resting membrane potential (*V*_rest_, fig.2c,d,g,h). The voltage-dependence of VSD-IV appeared shallower than VSD-III, indicating that VSD-IV is less sensitive to voltage changes. VCF mutations and labelling did not substantially change the voltage dependence of pore opening (fig.2f-h). Figure 2i illustrates the diverse conformational responses of different parts of α_1A_ to brief depolarizations.

**Fig. 2.**
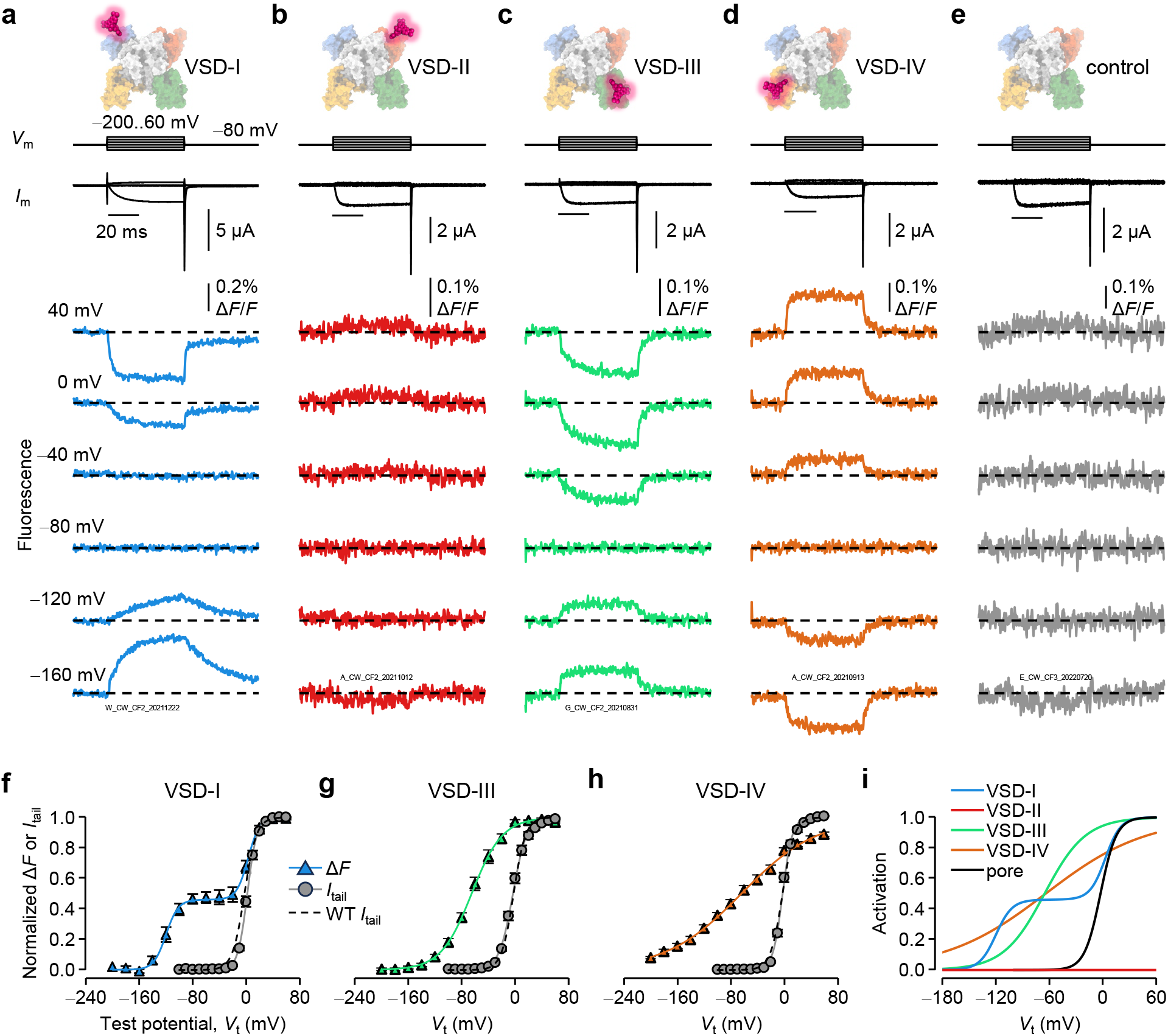
Ca_V_2.1 VSDs have diverse voltage-dependent-activation properties. **(a)** VCF recordings of Ca_V_2.1 complexes (α_1A_/α_2_δ-1/β_2a_) fluorescently labelled in VSD-I. Voltage steps (*V*_m_) are shown on top; ionic currents (*I*_m_) and fluorescence deflections (Δ*F*) were acquired simultaneously. **(b)** As in (a) for VSD-II. VSD-II does not show clear Δ*F* and appears to be voltage-insensitive (figs S1,S2). **(c**,**d)** As in (a) for VSD-III and VSD-IV, respectively. **(e)** As in (a) for control channels (no substituted Cys). **(f)** Voltage dependence of VSD-I activation (normalized Δ*F*, blue triangles) and fit to the sum of two Boltzmann distributions (blue curve, eq.3): *V*_0.5,*F1*_ = 3±1 mV, *z*_*F1*_ = 3.2±0.5 *e*_0_, *F1* = 52±4 %; *V*_0.5,*F*2_ =−119±3 mV, *z*_*F*2_ = 2.5±0.3 *e*_0_, *n* = 4 cells. Voltage dependence of pore opening (normalized *I*_tail_, eq.1) for VSD-I-labeled channels (grey circles and curve); *V*_0.5_ = 0.5±0.7 mV, *z*= 3.8±0.1 *e*_0_, *n* = 9. The voltage dependence of pore opening for control channels is shown as black dashed curve (*V*_0.5_ = −2.2±0.9 mV, *z* = 2.9±0.1 *e*_0_, *n* = 12). **(g)** As in (f), for VSD-III (green; eq.2; *V*_0.5_ = −65±4 mV, *z* = 1.1±0.1 *e*_0_, *n* = 7). Pore opening of VSD-III-labeled channels: *V*_0.5_ = −3±1 mV, *z* = 2.5±0.2 *e*_0_, *n* = 7. **(h)** As in (f), for VSD-IV (orange; VSD-IV *V*_0.5_ = −64±6 mV, *z* = 0.5±0.02 *e*_0_, *n* = 6). Pore opening of VSD-IV-labeled channels: *V*_0.5_ = −2±1 mV, *z* = 3.5±0.1 *e*_0_, *n* = 6. **(i)** Overlay of all voltage dependences observed on the human α_1A_ subunit. VSD-II activation is shown as a flat red line.

### Progressive VSD “conversion” under VDI-favouring conditions

Ca_V_2.1 availability is limited by VDI^10^ (fig.1a,b), here recapitulated by changing the holding membrane potential (*V*_h_; fig.3a). Which VSD is responsible for VDI? The VSD-I two-part response to membrane depolarization (fig.2f) suggested the presence of two Ca_V_2.1 populations: one whose VSD-I activated with similar voltage-dependence to pore opening and another whose VSD-I activated at very negative potentials, far (along the voltage axis) from pore opening. The latter process was reminiscent of “charge conversion”: the observations that charge movement (i.e., the overall activation of all VSDs measured by gating currents) occurs at more negative potentials as Ca_V_ channels enter inactivated states during prolonged depolarization^32, 33^. At negative *V*_h_ (−80 mV), a brief pulse to 40 mV produced robust VSD-I activation (fig.3b). In contrast, no VSD-I movements were detectable using the same step when *V*_h_ was very positive (40 mV, fig.3c), when channels were inactivated. In the presence of VDI-accelerating β_3_-subunits^25, 26^, fewer VSD-I could activate at *V*_h_ = −80 mV, compared with β_2a_ (fig.3b,d), and no movements were detected at *V*_h_ = 40 mV (fig.3e). These observations hinted that VSD-I is linked to VDI.

To explore *V*_h_-dependent VSD-I conversion in detail we used a broad range of *V*_h_ and brief (100-ms) test potentials (*V*_t_). Upon more positive *V*_h_, the proportion of channels with VSD-I with depolarized voltage-dependence (F1) progressively diminished (fig.3f; Boltzmann parameters in table S1), converting to channels whose VSD-I activated at hyperpolarized potentials (F2). A striking result was that F1 and F2 were separated by over 120 mV along the voltage axis, suggesting that conversion is a process that drastically alters the biophysical properties of VSD-I. Plots of the first derivatives of the voltage-dependence curves (fig.3g) and F1 percentage versus *V*_h_ (fig.3h) better illustrate F1-F2 interconversion, which occurred around *V*_rest_. In β_3_-containing channels, conversion to F2 was favoured, occurring at more negative *V*_h_ (fig.3i-k).

VSD-III and VSD-IV also converted (fig.4, table S1) and their apparently one-part voltage-dependences (fig.2g,h) could be reinterpreted as mixtures of two populations. The gap between F1 and F2 VSD-III activations was approximately half as wide as that of VSD-I (∼60 mV), suggesting that VSD-III is less altered by conversion. The F1 and F2 components of VSD-IV had strikingly different apparent voltage sensitivity. β_3_ facilitated VSD-III and VSD-IV conversion.

Summarizing our findings so far, the Ca_V_2.1 VSDs exhibit diverse responses to both transient depolarization (fig.2) and changes in the holding potential (figs.3,4).

**Fig. 3.**
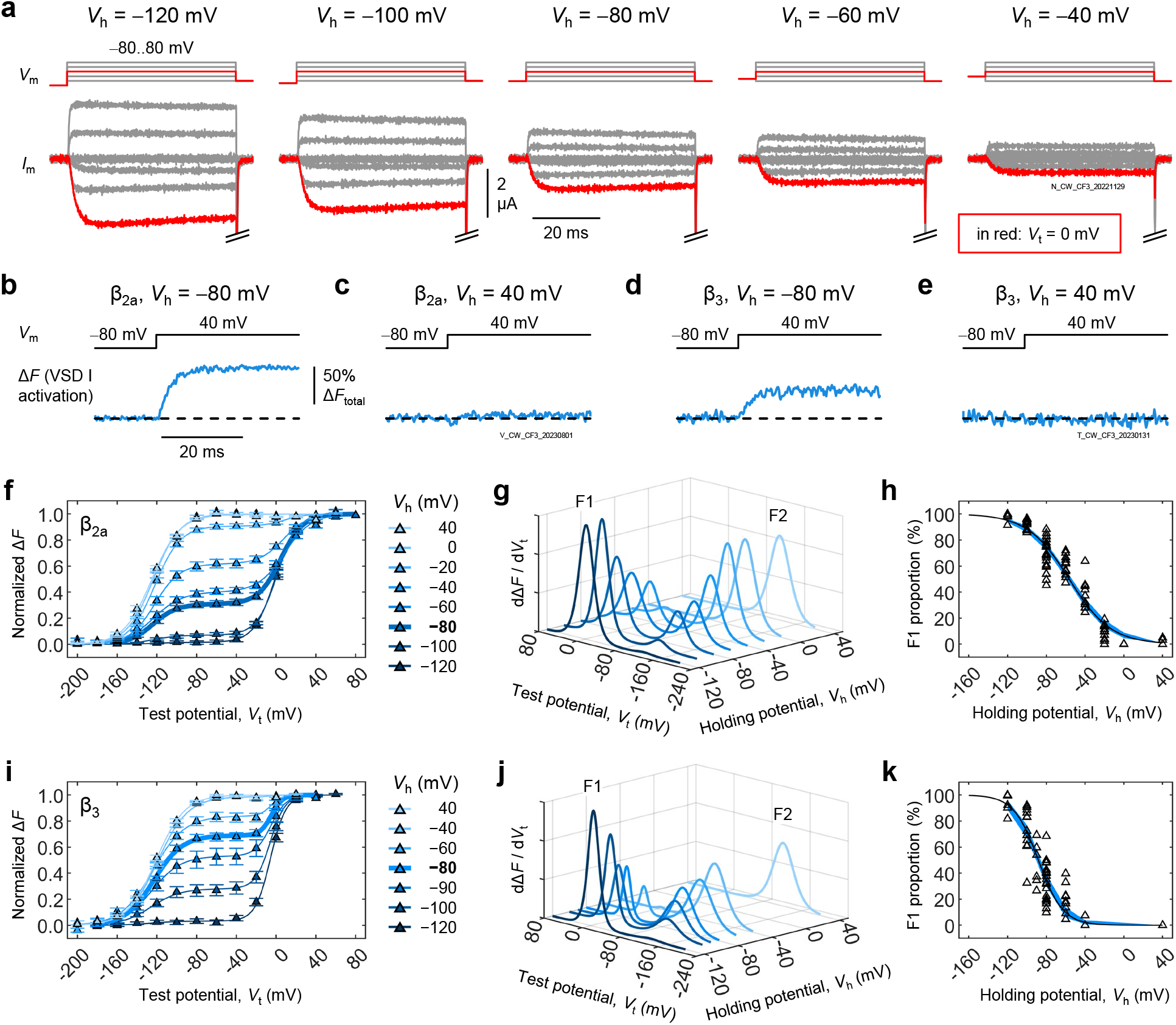
VSD-I converts under VDI-favouring conditions. **(a)** Voltage steps (*V*_m_) and exemplary currents (*I*_m_) from a cell expressing Ca_V_2.1 channels (α_1A_/α_2_δ-1/β_2a_) at different holding potentials (*V*_h_). Tail currents were cropped out for clarity. The current (i.e., channel availability) decreased as *V*_h_ became more positive: the hallmark of VDI^10^. **(b-e)** VSD-I activation in response to the same voltage step (−80 to 40 mV) under different VDI regimes: (b) β_2a_ subunits, *V*_h_ = −80 mV (VDI low); (c) β_2a_, *V*_h_ = 40 mV (VDI high); (d) β_3_, *V*_h_ = −80 mV (VDI intermediate); (e) β_3_, *V*_h_ = 40 mV (VDI high). The −80-mV steps in (c,e) were 100 ms long. **(f)** Voltage dependence of VSD-I activation at different *V*_h_ in the presence of β_2a_. Solid curves are the sums of two Boltzmann distributions (eq.3; parameters in table S1). Error bars are S.E.M. **(g)** The first derivatives of the curves from (f) illustrate the conversion of VSD-I from F1 to F2 as *V*_h_ becomes more positive. **(h)** Apparent voltage dependence of VSD-I conversion. Open triangles are individual data; the blue surface is the 95% confidence interval of a Boltzmann fit (eq.4; *V*_0.5_ = −56.4 [−59.0, −53.9] mV; *z* = 1.18 [1.05, 1.31] *e*_0_, *n* = 43 cells). **(i-k)** As in (f-h), respectively, for channels with β_3_. F1-F2 conversion occurs at more negative voltages: (*V*_0.5_ = −88.2 [−90.4, −86.0] mV; *z* = 2.00 [1.62, 2.37] *e*_0_, *n* = 23).

**Fig. 4.**
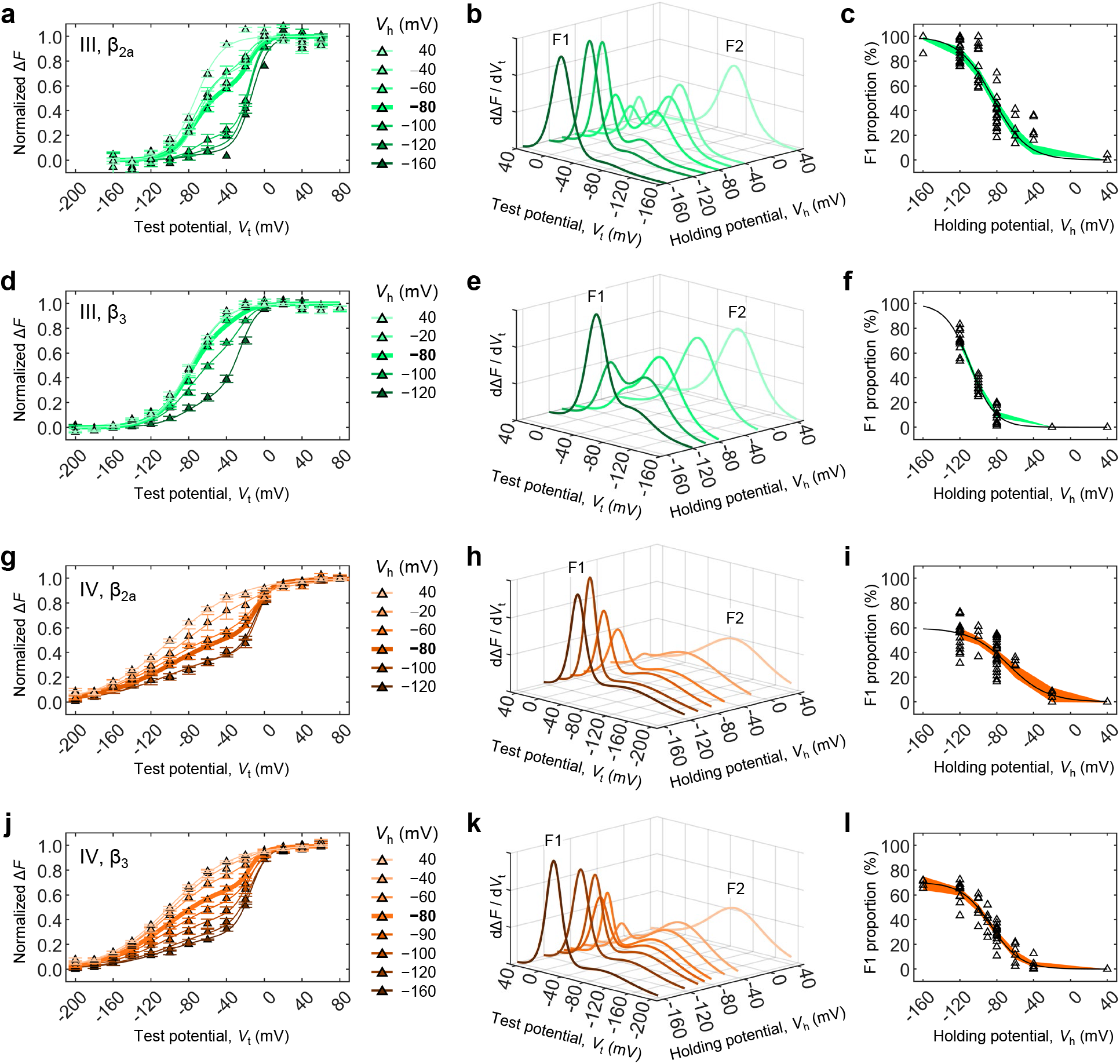
VSD-III and VSD-IV can convert. VSD-III and VSD-IV activation at extended holding and test potentials (*V*_h_, *V*_t_) revealed that they also undergo conversion. Similar to VSD-I, the voltage dependence of VSD-III and VSD-IV at *V*_h_ = −80 mV (fig.2g,h) consisted of transitions in a mixed population of “F1” and “F2”. **(a)** Voltage dependence of VSD-III activation in the presence of β_2a_. Solid curves are the sums of two Boltzmann distributions (eq.3, parameters in table S1). Error bars are S.E.M. **(b)** The first derivatives of the curves from (a) illustrate the conversion of VSD-III from “F1” to “F2” as *V*_h_ becomes more positive. **(c)** Apparent voltage-dependence of VSD-III conversion. Open triangles are individual data; the green surface is the 95% confidence interval of a Boltzmann fit (eq.4; *V*_0.5_ = −84.2 [−87.3,−81.1] mV; *z* = 1.37 [1.11,1.63] *e*_0_, *n* = 19 cells). **(d-f)** As in (a-c), respectively, for channels complexed with β_3_. The F1-F2 transition occurs at more negative voltages: (*V*_0.5_ = −109 [−110,−107] mV; *z* = 1.91 [1.69,2.12] *e*_0_, *n* = 13 cells). **(g-i)** As in (a-c), respectively, for channels with β_2a_ labelled in VSD-IV. In the Boltzmann fits of panel (i), the positive asymptote (F1_max_) was a free parameter: F1_max_ = 57.7 [49.9,65.5] %; *V*_0.5_ = −65.4 [−73.6,−57.2] mV; *z* = 1.28 [0.658,1.91] *e*_0_, *n* = 26 cells. **(j-l)** As in (g-i), respectively, for VSD-IV-labelled channels with β_3_. F1_max_ = 70.2 [64.0,76.3] %; *V*_0.5_ −86.3 [−90.5,−82.1] mV; *z* = 1.58 [1.20,1.97] *e*_0_, *n* = 15 cells.

### VSD-I conversion is linked to inactivation

Fitting fluorescence data to the sum of two Boltzmann functions provided a good empirical overview, but it had two shortcomings: (i) it implied that F1 and F2 transitions are independent, and (ii) it did not account for kinetics. To characterize VSD activation and conversion with more mechanistic rigor, VCF data from each VSD were fit to a four-state model (fig.5a). The kinetic model combines VSD activation and deactivation (responses to brief potential changes) as well as conversion between two modes of gating (responses to steady-state potential changes). Mode 1 corresponds to the F1 component from the Boltzmann-distribution fits, and mode 2 to F2. However, the four-state model is physically more meaningful, accounting for the kinetics of all transitions while obeying microscopic reversibility and charge conservation. The model fit to the data is shown in fig.S3, and the optimized parameters in table S2.

**Fig. 5.**
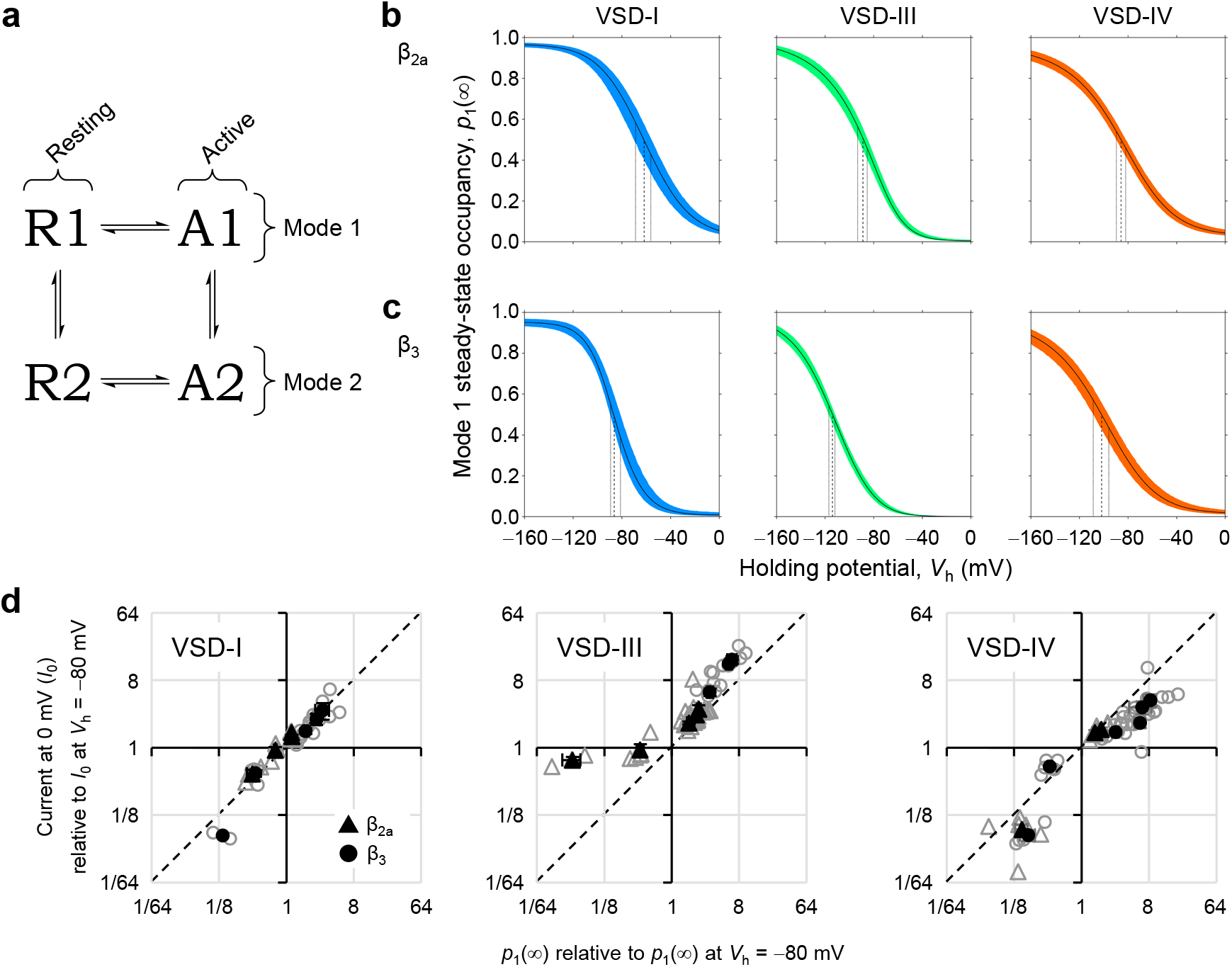
VSD-I conversion is linked to VDI. **(a)** The four-state model used to fit VCF data. Each VSD could achieve four conformations: R1: mode-1 resting state; A1: mode-1 active state; R2, A2: mode-2 resting and active states. Transitions are defined as: R→A: activation; A→R: deactivation; mode 1 → mode 2: conversion; mode 2 → mode 1: recovery. **(b)** The steady-state occupancy of mode-1 states (*p*_1_(∞), i.e., *p*_R1_(∞) + *p*_A1_(∞)) plotted against the holding potential (*V*_h_) in the presence of β_2a_. Colored area: 95% confidence interval; vertical dashed lines point to the mean *V*_0.5_; dotted lines: *V*_0.5_ 95% confidence intervals (parameters in table S2). **(c)** As in (b), now in the presence of VDI-favoring β_3_-subunits: all conversions are facilitated, occurring at more negative potentials (table S2). **(d)** Mode 1 occupancy (relative to Mode 1 at *V*_h_ = −80 mV) plotted against relative current availability. The latter was calculated from inward current at 0 mV (*I*_0_, as in the red traces in fig.3a) relative to *I*_0_ at *V*_h_ = −80 mV. Open symbols are from individual cells, filled symbols are means. VSD-I mode 1 occupancy was statistically indistinguishable from current availability (*p* = 0.967, *n* = 17 cells), in contrast to VSD-III (*p* = 0.0290, *n* = 26) and VSD-IV (*p* = 0.0226, *n* = 27). Kolmogorov-Smirnov two-sample tests. Error bars are S.E.M.

VSD-I converted from mode 1 to 2 “spontaneously”, i.e., in a voltage-independent manner. Conversion occured preferentially from state A1 (*k*_con_ = 0.43 s^−1^), while recovery occured from state R2 (*k*_rec_ = 0.17 s^−1^). Since A1 is visited at depolarized potentials, and R2 at hyperpolarized potentials, the distribution of channels in mode 1 or 2 had an apparent voltage dependence, with a *V*_0.5_ ≅ −60 mV (fig.5b). VSD-III and VSD-IV also converted spontaneously from the active state but, in contrast to VSD-I, the experimental fluorescence data were not consistent with a voltage-independent transition between R1 and R2. Instead, an intrinsically-voltage-dependent transition was required (*z*_R1↔R2_ = 0.98 and 0.56 *e*_0_, respectively; table S2). Their modal conversion occurred at more negative potentials (ca. −85 mV fig.5b, table S2) than for VSD-I.

Changing from β_2a_-to β_3_-subunits altered several biophysical parameters. Of note: First, the voltage dependence of VSD-I activation in mode 1 was accelerated by ∼15-fold and shifted to more negative voltages, separating from pore opening by ca. 25 mV (table S2). The activation transitions of other VSDs, and all activation transitions in mode 2, were relatively unaffected. Second, VSD-I A1→A2 conversion was accelerated by 8-fold, which resulted in a shift of the conversion voltage-dependence by −20 mV. Likewise, the conversions of VSD-III and VSD-IV were facilitated, resulting in similar negative shifts (fig.5b,c, table S2).

Most pertinent to VDI, the fraction of channels with a VSD-I in mode 1, and the fraction of channels available to activate (i.e., non-inactivated), were statistically indistinguishable. By contrast, the fraction of channels with VSD-III or VSD-IV in mode 1 were statistically distinct from the fraction of non-inactivated channels (fig.5d).

## Discussion

We have experimentally and analytically shown that the four Ca_V_2.1 VSDs display distinct conformational changes. The VSDs do not merely possess quantitatively distinct biophysical properties, but exhibit qualitative differences in their structural dynamics. VSD-I movements closely correlate with channel opening and VDI (figs.2f, 5d). VSD-II appears to be voltage-insensitive (figs.2, S1, S2). VSD-III and VSD-IV exhibit a voltage-dependent conversion from the resting state (table S2), which results in a steady-state occupancy of the mode-2 resting state (R2) over physiological *V*_rest_ (fig.6a,b): a unique feature, as R2 is a metastable state in “canonical”, spontaneously-converting VSDs^34, 35^, like VSD-I. To better illustrate the multiplicity of VSD steady-state conformations, we mapped the state occupancies of each VSD into “state spectra” (fig.6a,b), used to color the Ca_V_2.1 structure (fig.6c, movie S1).

**Fig. 6.**
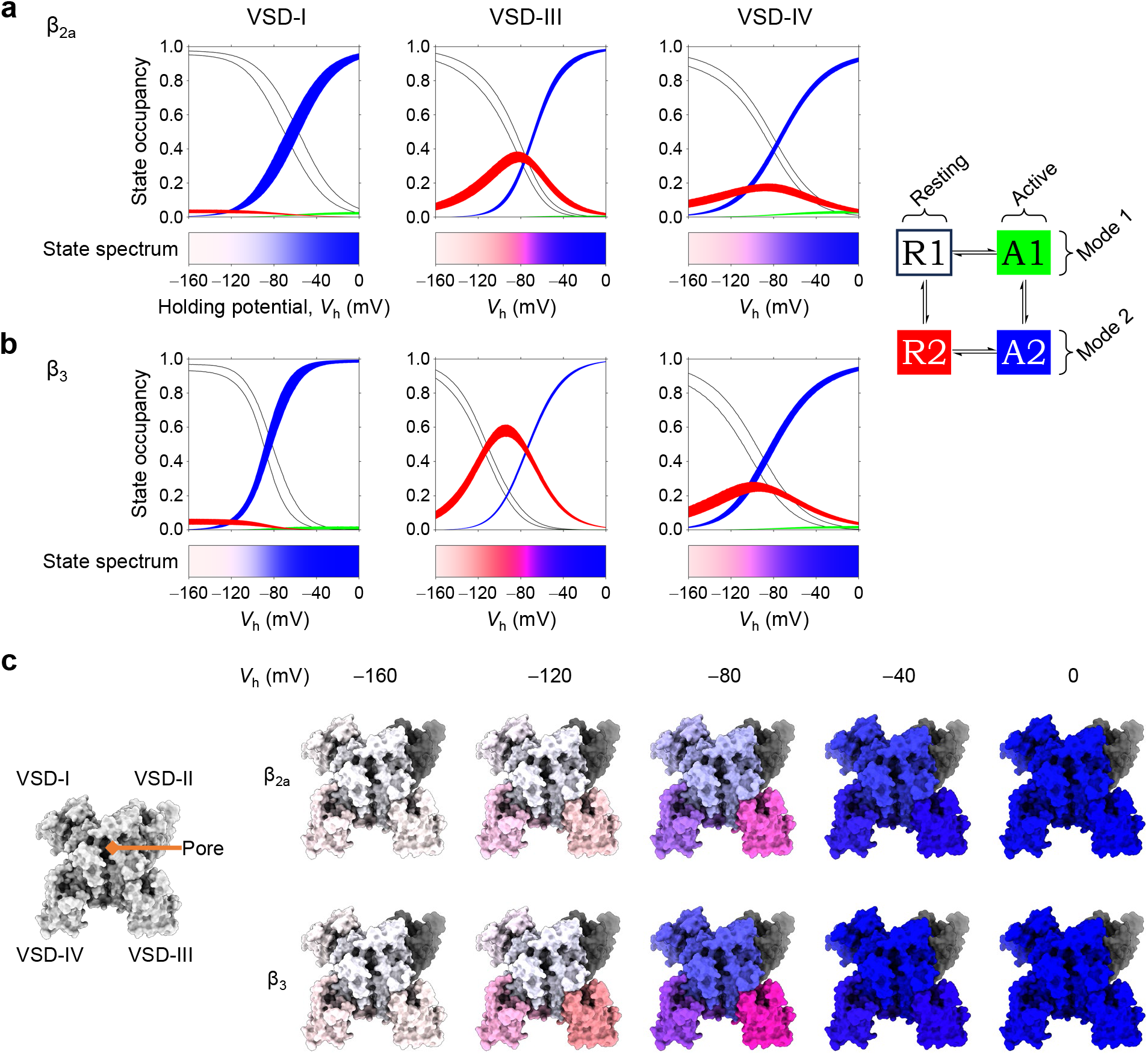
The rich conformational palette of Ca_V_2.1 at equilibrium. **(a)** Steady-state curves of all states (color code on the scheme on the right), plotted against the holding potential (*V*_h_). Areas are 95% confidence intervals. VSD-I behaves as a canonical converting VSD, with states A1 and R2 being metastable (very low steady-state occupancy). The voltage-dependent R1-R2 conversions in VSD-III and VSD-IV result in stable R2 states around physiological resting potentials. The “state spectra” colorbars below encode the state occupancies into color information. **(b)** As in (a), for channels complexed with β_3_. The VDI-favoring subunit alters the overall state spectra for all VSDs. **(c)** Color information in the state spectra was used to annotate the α_1A_ surface (PDB: 8X90^31^) at different *V*_h_ and β-subunits. The white-to-blue transitions illustrate the R1-to-A2 modal shift in VSD-I, while VSD-III and VSD-IV exhibit prominent red/purple hues, due to the stable occupancy of R2 states. The pore is colored white for closed, blue for inactivated; as inactivation best correlated with VSD-I modal shifts (fig.5d), it follows the same color. VSD-II is shown in grey. Movie S1 is an animated version of this figure.

A major finding is the relevance of VSD-I transitions to Ca_V_2.1 opening and VDI. VSD-I activation (in mode 1) and pore opening occur over the same membrane potential (fig.2, table S2). We propose that such processes may be called *syntasic*, from classical Greek syn (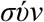, together) and tasis (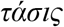, tension, or in this case, voltage). Effectively, VSD-I activation to A1 is the first molecular transition that triggers neurotransmitter release in most synapses.

VSD-I conversion (shift to mode 2), tuned to *V*_rest_, is linked to VDI (fig.5). That is, *V*_rest_ bisects the Ca_V_2.1 population into channels with VSD-I in mode 1, primed to trigger neurosecretion; and channels with VSD-I in mode 2 and inactivated, but available to be recruited when *V*_rest_ becomes more negative. A straightforward mechanism for this is that, since VSD-I A1 is linked to pore opening, inability to achieve A1 would produce channels unavailable to conduct. VSD-I conversion may, as a conformational change, also play an active role in VDI development, engaging cytosolic structures lacking intrinsic voltage dependence yet associated with inactivation, such as the hinged lid^36^ and the W-helix^37, 38, 39^. Since Ca_V_2.1 VDI contributes to synaptic plasticity mechanisms^10, 11, 14, 15^, another VSD-I transition—in this case conversion—is shown to be linked to processes of a scale well beyond intramolecular structural dynamics: cognition, and memory formation. Recovery from conversion takes several seconds (*k*_rec_ = 0.17 or 0.30 s^−1^ with β_2a_ or β_3_, respectively; table S2), by far exceeding the near-millisecond duration of a neuronal spike. This demonstrates how conversion acts as a memory mechanism in this channel. Moreover, every single spike has a small probability of triggering VSD-I conversion. The onset of VSD-I conversion is a relatively slow transition for an ion-channel molecule, with kinetics in the order of a second (*k*_con_ = 0.43 or 3.3 s^−1^ with β_2a_ or β_3_, respectively; table S2); and yet, as a conformational change that could culminate in acquiring lifelong memories, it is also one of the fastest events on the “timescales of learning”^40^.

The lack of VSD-II voltage-dependence, both in Ca_V_2.1 (figs.2, S1, S2) and Ca_V_2.2^41^ is a consistent feature. An inability to undergo voltage-dependent movements explains why, in all Ca_V_2-channel structures reported, VSD-II was resolved in a resting conformation^31, 37, 38, 39, 42^ despite the absence of an electric field (equivalent to *V*_h_ = 0 mV). In these structures, all other VSDs were resolved in an active state. By the same token, VSD-II being “locked down” in the resolved Ca_V_2 structures supports the lack of optical signals reported here and for CaV2.2^41^. Whether VSD-II can activate under a different set of conditions is an outstanding question.

Both VSD-III and VSD-IV activate faster than VSD-I and at more negative potentials (table S2); they could possess regulatory roles in the activation process. In β_3_-containing channels, where VSD-I activation and pore opening are “asyntasic” (*V*_0.5_ of −24 and 5 mV, respectively; table S2), voltage-dependent opening could be a more cooperative process involving VSD-III and VSD-IV. A plasticity of VSD-pore connectivity following auxiliary-subunit changes has been reported in Ca_V_1.2^43^. A common feature of Ca_V_-channel VSDs is that, despite their homology, their non-identity translates to functional heterogeneity—there are functional differences both within and between different Ca_V_ isoforms^17, 41, 44^.

β_2a_, relative to β_3_, strongly inhibited modal shifts by shifting the overall steady-state conversion to more positive potentials (figs 3-5, table S2). VSD-I and VSD-III are more affected by changes in β-subunit composition than VSD-IV. The most pronounced effects of β_2a_ are a slower conversion from A1 to A2 in VSD-I (by ∼8-fold; table S2) and an acceleration of the recovery from R2 to R1 in VSD-III (by ∼7-fold at −80 mV; calculated from parameter values in table S2 using eq.8). β-subunits bind to the cytosolic I-II loop^36, 45^ (fig.1a). While we cannot exclude allosteric effects of β-subunits on VSD structural dynamics, β-subunits may also interact directly with the cytosolic VSD flanks, in a state-dependent manner, similar to the proposed action of G-proteins on Ca_V_2.2^41^.

Our work here has uncovered a particularly rich gamut of conformations of the Ca_V_2.1 VSDs, as they respond to electrical signals both transient and long-lived, tuned to both the resting membrane potential and depolarization, and under the influence of different β-subunits. Yet this is only a part of the regulation Ca_V_2.1 are subject to in their presynaptic environment: several molecular partners, including Gβγ^46, 47^, calmodulin^48^ and CaBP1^49^, as well as neuronal junctophilins^50^ and syntaxin^51^, can modulate Ca_V_2.1 voltage-dependent activation and inactivation. It will be of high interest to investigate whether they act via the same pathway as β-subunits, or whether the Ca_V_2.1 voltage-sensing apparatus possesses specific handles for each regulatory partner. Given the importance of Ca^2+^-signal amplitude and timing for synaptic communication, it is fitting that its principal mediator is a macromolecule with exquisite structural dynamics and regulation.

## Methods

### Molecular biology

The human *CACNA1A* transcript variant 3 (EFa, NM_001127221.2, Uniprot O00555.3) was codon optimized for *Xenopus laevis* expression by Integrated DNA Technologies (IDT) and subcloned into the Z-vector^52^. All site-directed mutagenesis was performed with a high-fidelity *Pfu* polymerase (Agilent 600850) and confirmed by full-gene DNA sequencing. Molecular biology reagents were obtained from New England Biolabs, and synthetic oligonucleotides from IDT. *In vitro* cRNA transcription was performed with the AmpliCap-Max T7 High Yield Message Maker Kit (Cellscript); RNA was stored at −80 ^o^C in RNA storage solution (Thermo Fisher Scientific). α_1A_ subunits were coexpressed with rabbit α_2_δ-1 (*CACNA2D1*, Uniprot P13806) and either rat β_2a_ (*CACNB2A*, UniProt Q8VGC3) or rabbit β_3_ (*CACNB3*, Uniprot P54286) subunits.

### Oocyte preparation and labelling

All animal experiments were approved by the Linköping University Animal Care and Use Committee (document number 15839-2018, protocol number 1941). Defolliculated *Xenopus laevis* (Nasco) oocytes (stage V-VI) were prepared as previously described^41^ or purchased from Ecocyte. Each oocyte was micro-injected with a 50 nL cRNA mixture of α_1A_, α_2_δ-1, and either β_2a_ or β_3_ (0.6-0.8 μg/μL of each subunit). Oocytes were incubated at 17 ^o^C in 0.5× Leibovitz’s L-15 (Corning) diluted in MilliQ H_2_O, supplemented with 1% horse serum (Capricorn Scientific), 100 units/mL penicillin and 100 μg/mL streptomycin (Gibco), 100 μg/mL amikacin (Fisher BioReagents) for 4-6 days. Prior to fluorescence staining, oocytes were rinsed in SOS (in mM: 100 NaCl, 2 KCl, 1.8 CaCl_2_, 1 MgCl_2_, 5 HEPES; pH=7.0).

Oocytes expressing Cys-substituted Ca_V_2.1 channel complexes were labelled with, unless otherwise stated, 20 μM MTS-5(6)-carboxytetramethylrhodamine (MTS-TAMRA; Biotium) for 7 minutes at 4 ^o^C in a depolarizing solution (in mM: 120 K-Methanesulfonate (MES), 2 Ca(MES)_2_, 10 HEPES; pH=7.0). Alternate fluorophores attempted for VSD-II were: 10 μM tetramethylrhodamine-6-maleimide (TMR6M; AAT Bioquest) for 15 minutes at 4 ^o^C, 20 μM tetramethylrhodamine-6-maleimide C6 (6-TAMRA C6 maleimide; AAT Bioquest) for 25 minutes at room temperature, 100 μM Alexa Fluor 488 C_5_ maleimide (Alexa-488; Thermo Fisher Scientific) for 30 minutes on ice. Oocytes were rinsed in dye-free SOS following fluorescence labelling.

### Electrophysiological techniques

Oocytes were voltage-clamped under the cut-open oocyte Vaseline Gap (COVG) technique complemented with epifluorescence detection^20, 22, 41^. A CA-1B amplifier (Dagan Corporation) was used in COVG mode. Data were acquired at 25 kHz using a Digidata 1550B1 digitizer and pClamp 11.2.1 software (Molecular Devices). The optical set-up consisted of a BX51WI upright microscope (Olympus) with filters (Semrock BrightLine: exciter: FF01-531/40-25; dichroic: FF562-Di02-25×36; emitter: FF01-593/40-25). The excitation light source was the M530L3 green LED (530 nm, 170 mW, Thorlabs) driven by a Cyclops LED driver (Open Ephys). For Alexa-488 experiments, the following filter set was used (Semrock BrightLine): exciter: FF01-482/35-25; dichroic: FF506-DI03-25×36; emitter: FF01-524/24-25. The light source was a Thorlabs blue LED (490 nm, 205 mW, M490L4). A LUMPLANFL 40XW water immersion objective (Olympus; numerical aperture = 0.8, working distance = 3.3 mm) and SM05PD3A Si photodiode (Thorlabs) were used for fluorescence detection. Photocurrent was amplified with a DLPCA-200 current amplifier (FEMTO). Fluorescence emission and ionic currents were simultaneously recorded from the oocyte membrane isolated by the top chamber and low-pass-filtered at 5 kHz.

Prior to recordings, oocytes were injected with 100 nL of 100 mM BAPTA*•*4K, 10 mM HEPES, pH=7.0 to prevent activation of endogenous Ca^2+^-and Ba^2+^-dependent Cl^−^ channels . External solution (in mM): 120 NaMES, 2 Ba(MES)_2_, 10 HEPES; pH=7.0. Internal solution (in mM): 120 K-Glutamate, 10 HEPES; pH=7.0. Intracellular micropipette solution: 3 M NaMES, 10 mM NaCl, 10 mM HEPES; pH=7.0. Oocytes were permeabilized using 0.1% saponin to gain low resistance intracellular access. Unless otherwise stated, oocytes were clamped at a holding potential of −80 mV. To evaluate the voltage dependence of channel activation, a series of 50 ms test pulses from −100 mV to 80 mV, in 10 mV increments, was used. P/−6 subtraction was performed to reduce capacitive transients. To examine the voltage dependence of VSD-III and VSD-IV activation, 50 ms test pulses within the range of −200 mV to 60 mV, in 20 mV increments, was used. For VSD-I, an activating pulse of 100 ms was used unless otherwise stated, as fluorescence deflections did not achieve steady-state by 50 ms. 4 averages were performed to increase the signal-to-noise ratio of fluorescence signals. To evaluate different holding potentials (*V*_h_), oocytes were clamped to each *V*_h_ for 2 min to allow complete equilibration/conversion of channels prior to running experimental protocols.

### Data analysis

The voltage dependence of channel opening was obtained from the peak tail current at *V*_h_=−80 mV and fit to the single Boltzmann function:

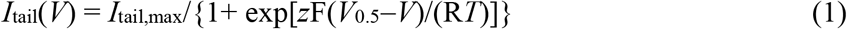

where *V* was membrane potential, *I*_tail,max_ was the maximal *I*_tail_, *z* was the valence, *V*_0.5_ was the half-activation potential, F was the Faraday constant, R was the gas constant and *T* was temperature (294 K).

The voltage dependence of fluorescence deflection (Δ*F*) was obtained from the average fluorescence signal during the last 5 ms of the test pulse. Δ*F* for VSDs III and IV were fit to the single Boltzmann function:

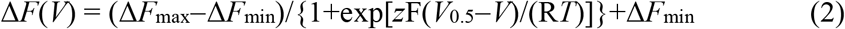

where Δ*F*_max_ and Δ*F*_min_ were the maximal and minimal Δ*F* asymptotes, respectively. In the case of the Δ*F*(*V*) curve for VSD-I (fig.2f), and subsequent fittings of VSDs I, III and IV at extended holding potentials (figs.3f,i & 4a,d,g,j), the sum of two Boltzmann distributions was used:

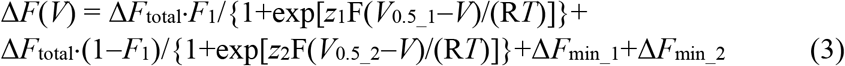

where Δ*F*_total_ was the total fluorescence change (Δ*F*_max_1_+Δ*F*_max_2_−Δ*F*_min_1_−Δ*F*_min_2_) and *F*_1_ was the fractional amplitude of the depolarized fluorescence component [(Δ*F*_max_1_−Δ*F*_min_1_)/Δ*F*_total_)] To help define the parameters, only cells with >1 *V*_h_ were fit, and the following constraints were placed to reduce the number of free parameters:

1) Voltage-dependence parameters (*V*_0.5_1_, *z*_1_, *V*_0.5_2_ and *z*_2_) were constrained to be equal across fits of different *V*_h_ for each cell.
2) Δ*F*_min_1_ was constrained to be equal to Δ*F*_max_2_ for each *V*_h_, in each cell.

Fitting was performed by least squares using *Solver* in Microsoft Excel. Data are represented as mean ± S.E.M.

To determine the apparent voltage-dependence of VSD conversion from the Boltzmann fits (figs 3h,k & 4c,f,i,l), *F*_1_ values from all cells and *V*_h_ were pooled together and fit to the Boltzmann distribution:

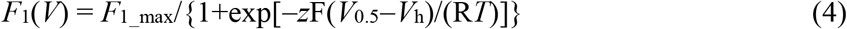

*F*_1_max_ was fixed to 1 for VSD-I and VSD-III, and left as a free parameter for VSD-IV. Fitting was perfomed in Mathworks Matlab using *fit*. 95% confidence intervals were estimated using Mathworks Matlab *confint*.

A four-state model was constructed in Matlab R2019a (MathWorks) representing transitions between a VSD active and resting states between modes 1 and 2. Activation transition rates (A1→R1 and A2→R2) were modelled as:

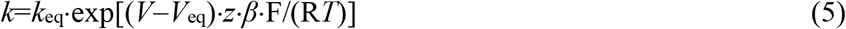

while deactivation transition rates (A1←R1 and A2←R2) as:

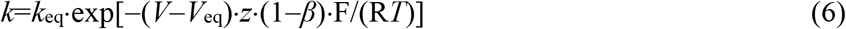

where *k*_eq_, *V*_eq_, *z* and *β* are shared free parameters. *k*_eq_ is an equilibrium rate constant, *V*_eq_ is the equilibrium potential, and *β* is the portion of position of the energy barrier on the electric field. Conversion rate (R1→R2) was modelled as:

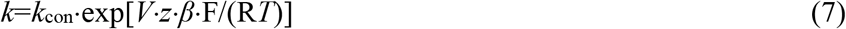

while conversion recovery rates (R1←R2 and A1←A2) were modelled as:

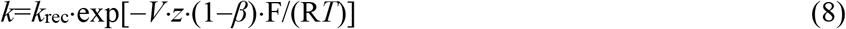

Each conversion/recovery equilibrium also shared four free parameters (*k*_con_, *k*_rec_, *z, β*), but this formulation was more easily adaptable to becoming voltage-independent, by fixing *z* to 0.

To obey microscopic reversibility, conversion rate (A1→A2) was calculated by:

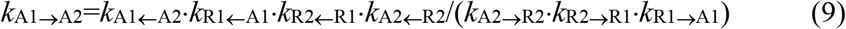

This maneuver also reduced the number of free parameters by one, as *k*_con_ did not have to be calculated for the A1↔A2 equilibrium.

Finally, to obey conservation of charge, the valence of the R2↔A2 equilibrium was also excluded as a free parameter, and was calculated by:

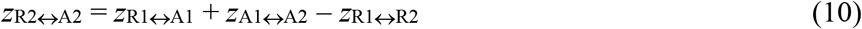

The model rates were formulated into a **Q**-matrix^53, 54^. Briefly, **Q** was a square 4×4 matrix. Each element *q*_*i*,*j*_ contained the rate for the transition from state *i* to state *j*. If there were no connection between states *i* and *j* then *q*_*i*,*j*_=0. Each diagonal element was the negative sum of the off-diagonal elements in its row. In this way,

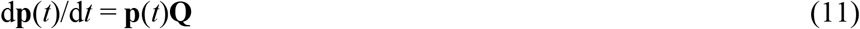

where **p**(*t*) was a 1×4 vector of probability (occupancy) for each state. The Matlab *ode15s* solver was used to calculate it. The voltage steps had a 43-μs time-constant to both emulate the COVG clamp speed and reduce stiffness. For initial conditions, background fluorescence calculations, and other calculations after fitting, the state occupancies at steady-state were calculated using:

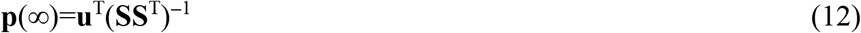

where **u** was a 4×1 unitary vector and **S** was [**Q u**].

Finally, 4×1 vector **f** contained fluorescence levels of each state. State R1 fluorescence was fixed to 0. Fluorescence was simulated as:

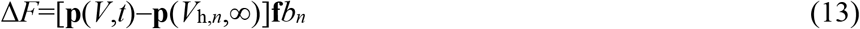

where *V*_h,*n*_ is the holding potential of the *n*th recording from the cell, and *b*_*n*_ was a factor to account for fluorescence bleaching during the experiments, which reduced the Δ*F* amplitude. *b*_1_ was fixed to 1, and *b*_*n*>1_ had bounds 0 and 1.

Data from each cell with >2 *V*_h_ were fit simultaneously. Rate optimization was performed by least squares, using the Bayesian adaptive direct search (BADS) machine-learning, model-fitting algorithm^55^.

The formulae for the R1-R2 rates did not contain an equilibrium-potential parameter (eqs.6,7). When the R1-R2 equilibrium was voltage-dependent (*z* > 0, for VSD-III and VSD-IV), *V*_eq_ was calculated after fitting using:

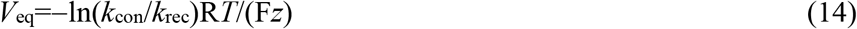

Finally, the equilibrium potential of modal shift was calculated iteratively using Matlab’s *lsqcurvefit*, solving for the voltage where sum of the mode-1 steady-state occupancies was 0.5. After several cells were fit, optimized and calculated parameters were averaged: the geometric mean was used for rate-constant parameters (*k*_eq_, *k*_con_, *k*_rec_), while the arithmetic mean was used for all others. 95% confidence intervals were calculated by bootstrapping (Matlab *bootci*, 10000 iterations).

Mode-1 occupancy and available current correlations (fig.5d) were performed as follows: Available current was calculated using test-pulses to 0 mV with different *V*_h_, which produced inward current according to channel availability (red traces in fig.3a). For each cell, the currents measured were normalized to the current with *V*_h_ = −80mV, which was available in all recordings. Only cells whose Δ*F* were fit with the 4-state model were included in this dataset. Mode-1 occupancy was calculated as the sum of occupancies of states R1 and A1 using eq.12, and normalized to mode-1 occupancy with *V*_h_ = −80 mV. Two-sample Kolmogorov-Smirnov tests were used to compare the distributions of available channels and channels in mode 1.

State occupancies were converted into color information (“state spectra”, fig.6a,b) by first assigning the occupancies of states R2, A1 and A2 as red-green-blue (RGB, respectively) triplets. To encode the fourth state (R1) occupancy, the RGB triplets were converted into the hue-saturation-lightness (HSL) color model. The lightness values were then replaced by:

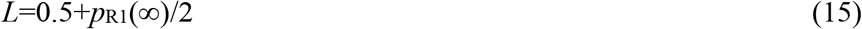

where *p*_R1_(∞) is the steady-state occupancy of R1. In this way, when *p*_R1_(∞) = 0, the spectrum has medium lightness, allowing the underlying color to show; when *p*_R1_(∞) = 1, the spectrum has maximal lightness (white). The HSL triplets were then converted back into the RGB format, to construct the spectra or annotate the Ca_V_2.1 structure (fig.6 & movie S1).

### Protein structure rendering

Structures of the human Ca_V_2.1 α_1A_ subunit (PDB: 8X90^31^) were rendered on UCSF ChimeraX^56, 57, 58^ and PyMOL (Schrödinger).

## Acknowledgments

We thank members of the Pantazis group for useful discussions and members of the Elinder, Liin and Pantazis groups for oocyte preparation. Funding: Lions Forskningsfond Ph.D. support (M.N.), NIH/NIGMS R35GM131896 (R.O.), start-up funds from the Linköping University Wallenberg Center for Molecular Medicine / the Knut and Alice Wallenberg Foundation (A.P.) ,Hjärnfonden (The Swedish Brain Foundation) grants and FO2022-0219 (F.E.), FO2022-0003 and FO2023-0025 (A.P.), Hjärt-Lung Fonden (The Swedish Heart-Lung Foundation) 20210596 (F.E.), Vetenskapsrådet (The Swedish Research Council) grants 2020-01019 (F.E.), and 2019-00988 and 2022-00574 (A.P.).

## Author contributions

K.W., M.N and M.A performed experiments. K.W., M.N. and A.P. performed analysis. R.O. and F.E. contributed reagents and methods. K.W., M.N., F.E. and A.P. wrote the manuscript. All authors contributed to manuscript review and editing.

## Competing interests

Authors have no competing interests.

**Fig. S1.**
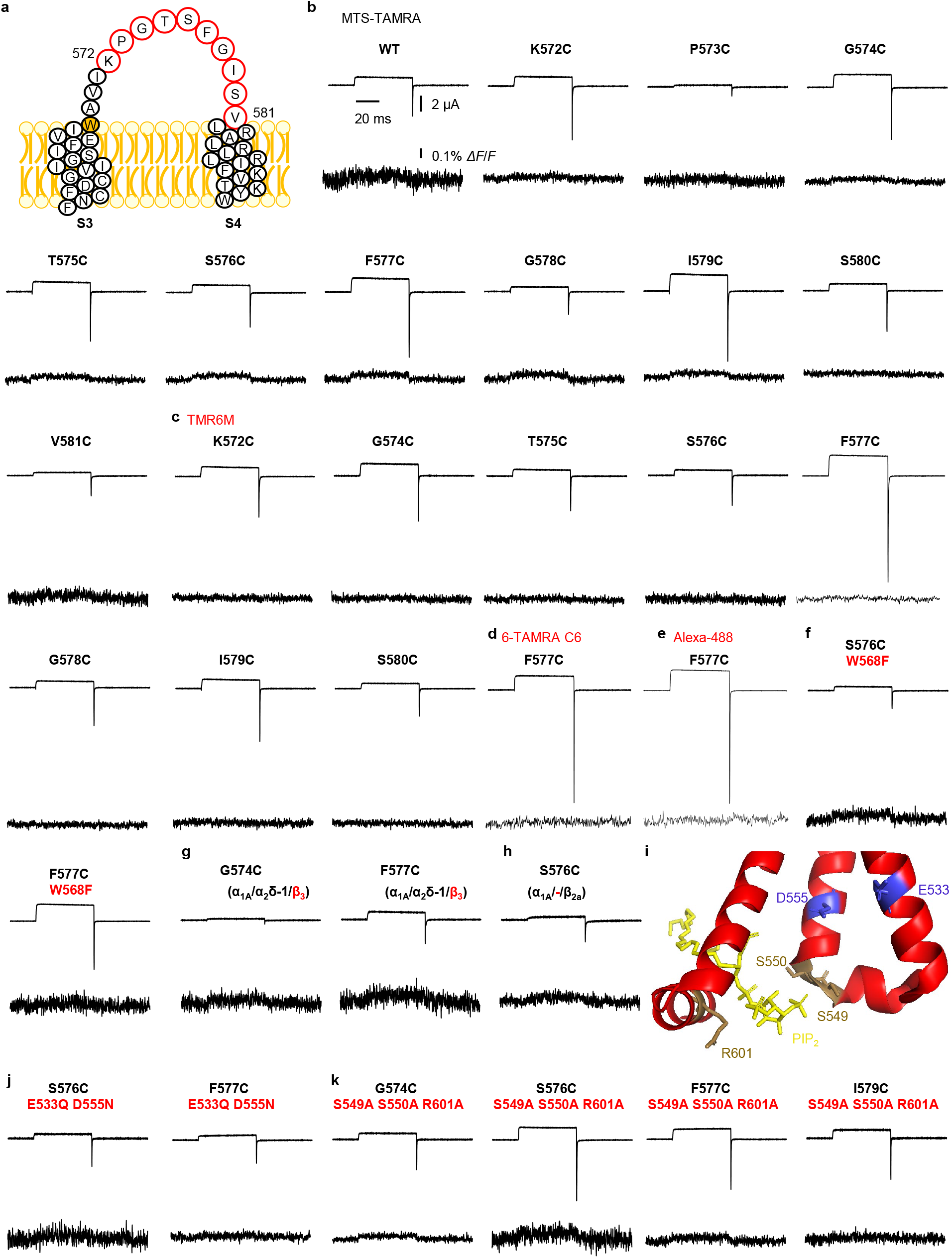
No voltage-dependent activation of Ca_V_2.1 VSD-II is detected. **(a)** Snake plot of the S3-S4 segment, and extracellular linker, of Ca_V_2.1 VSD-II. Positions tested by VCF are indicated by red circles. **(b)** Current and fluorescence traces from MTS-TAMRA-labelled Ca_V_2.1 channel complexes (α_1A_ construct indicated + α_2_δ-1 + β_2a_) in response to a voltage step from −80 mV to 80 mV. **(c-e)** As in (b) for channels labeled with fluorophores TMR6M, 6-TAMRA C6 maleimide or Alexa-488 maleimide, respectively. **(f-h)** As in (b), using channels with a substituted Trp in S3, co-expressed with the β_3_ subunit, or lacking the α_2_δ-1 subunit, respectively. **(i)** Magnified view of the PIP_2_-binding site in VSD-II (PDB: 8X90^31^). PIP_2_-binding residues (sand-coloured) and counter-charge residues (blue) are indicated. **(j**,**k)** As in (b), using channels lacking two counter-charge residues in S2-S3, or the PIP_2_ binding residues, respectively. Despite extensive efforts, no fluorescence deflections are observed from Ca_V_2.1 channels fluorescently labelled in VSD-II in response to membrane depolarization.

**Fig. S2.**
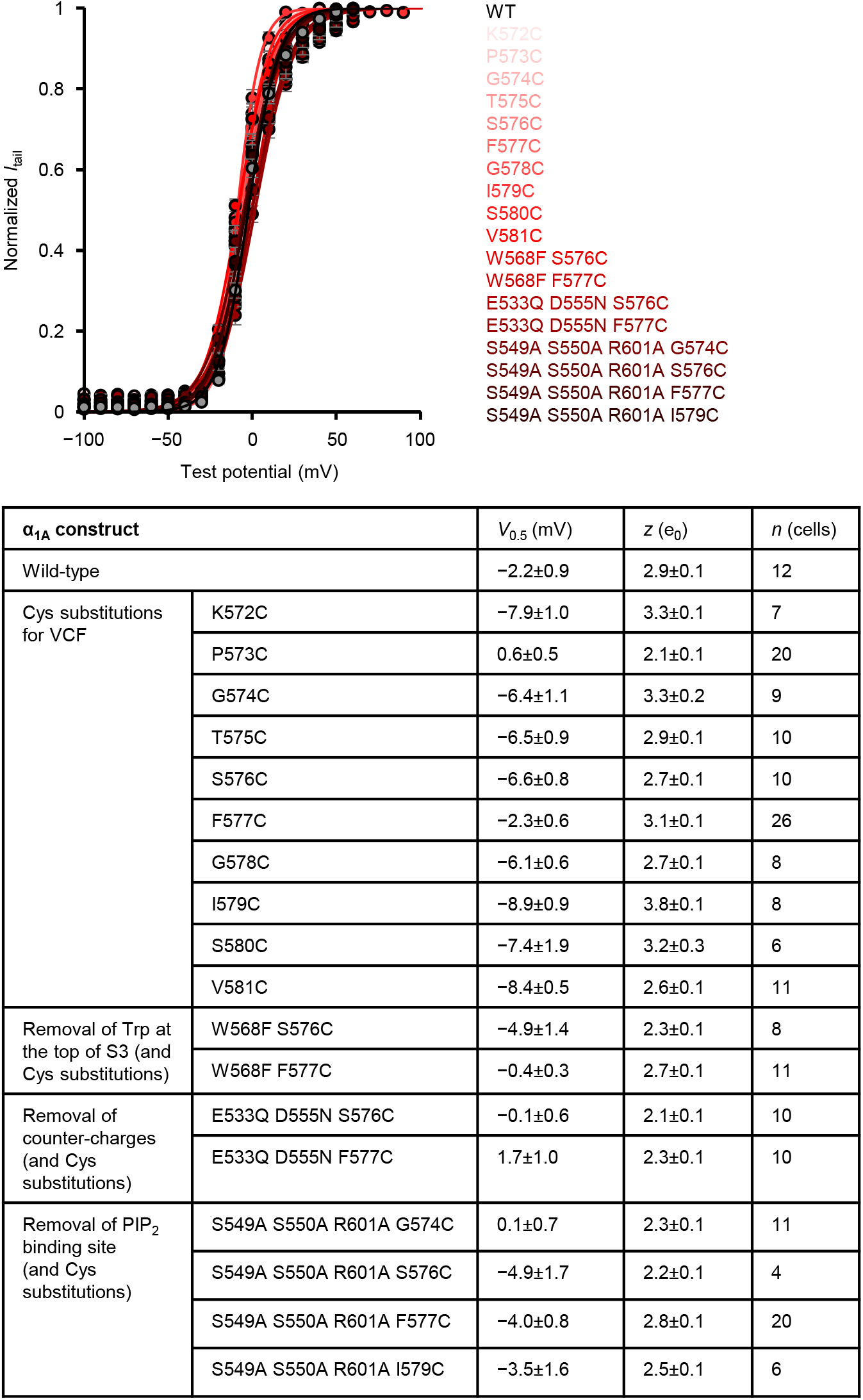
Mutations in VSD-II have minimal effects on Ca_V_2.1 voltage-dependent opening. Normalized tail current-voltage relationships were fit to a Boltzmann distribution (eq.1). Error bars represent S.E.M. Introduction of point mutations in VSD-II (and fluorescent labelling with MTS-TAMRA) did not substantially alter the voltage-dependence of pore opening compared to wild-type channels.

**Fig. S3.**
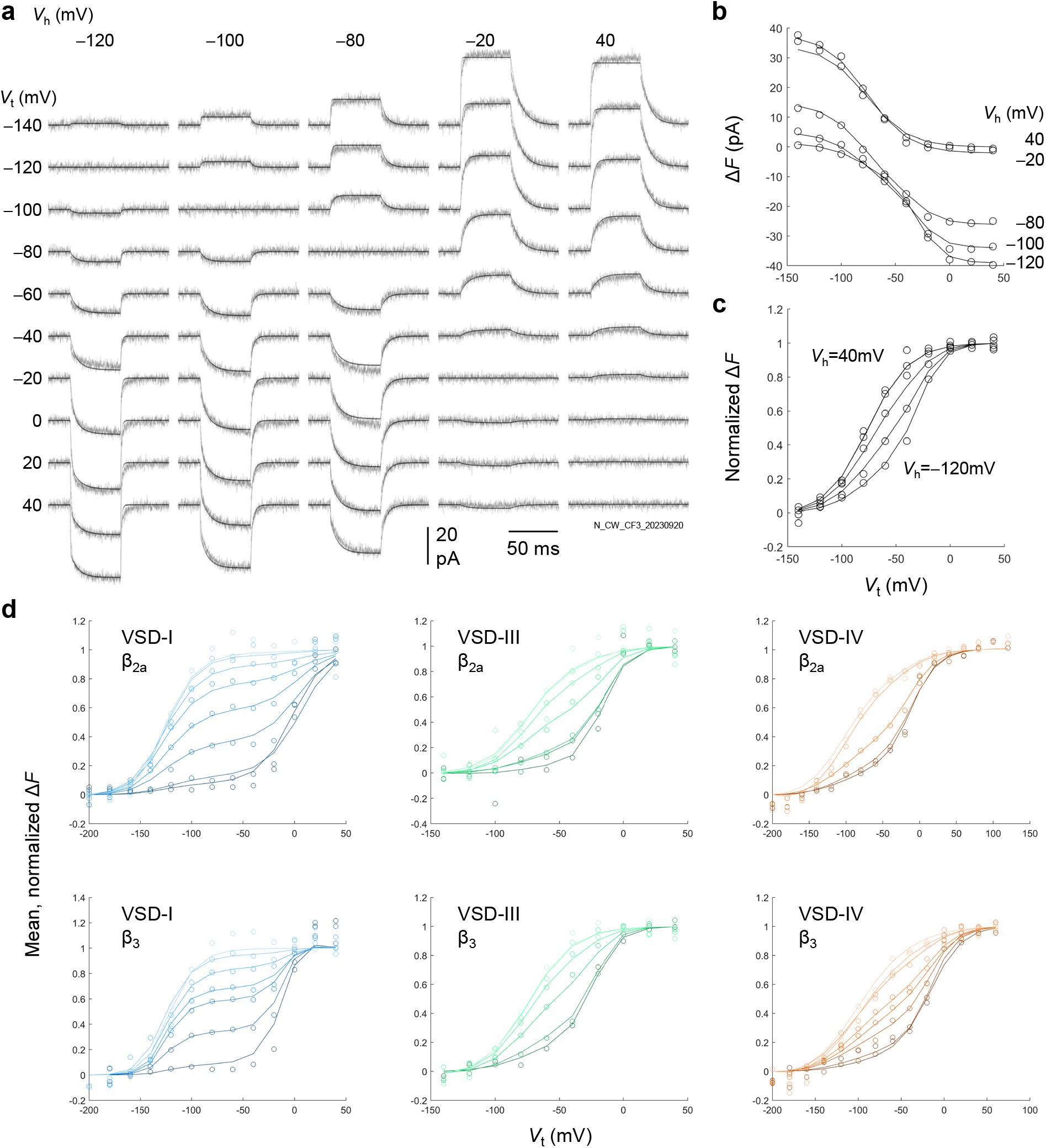
4-state model fitting. **(a)** Fitting of Δ*F* traces in a representative cell expressing VSD-III-labeled Ca_V_2.1-channels with β_3_. VSD-III activation was probed with test potentials (*V*_t_) from −140 to 40 mV over the holding potentials (*V*_h_) indicated, from −120 to 40 mV. Grey traces are the Δ*F* data; black lines are the model output (eq.13). A total of 14 cells from this condition (VSD-III, β_3_) were fit thus, and 81 cells across all conditions. **(b**,**c)** Model fits (black curves) over the raw (b) and normalized (a) Δ*F* (open circles) from the cell in (a). **(d)** Mean model fits (curves) and mean, normalized Δ*F* (open circles). Parameters are in table S2.

**Table. S1.** Conversion Boltzmann parameters. Equation 3 was used. Errors are S.E.M.

|  |  |  |  |  |  |  |  |  |  |
| --- | --- | --- | --- | --- | --- | --- | --- | --- | --- |
| VSD-I<br>$\beta_{2a}$<br>(Fig.3f) | $V_h$ (mV) | -120 | -100 | -80 | -60 | -40 | -20 | 0 | 40 |
| | $F_1$ (%) | 98.5 ± 1.04 | 92.3 ± 1.23 | 69.2 ± 2.37 | 60.0 ± 2.21 | 38.3 ± 3.45 | 9.66 ± 1.75 | 0.0175 ± 0.0175 | 0.489 ± 0.343 |
| | $V_{0.5,1}$ (mV) | -0.996 ± 1.09 | 2.62 ± 1.56 | 4.31 ± 0.966 | 7.02 ± 1.54 | 7.39 ± 0.951 | 8.00 ± 0.899 | 5.18 ± 1.96 | 6.85 ± 1.24 |
| | $z_1$ ($e_0$) | 2.40 ± 0.157 | 2.53 ± 0.138 | 2.36 ± 0.103 | 2.11 ± 0.0987 | 2.12 ± 0.124 | 2.10 ± 0.124 | 1.87 ± 0.185 | 1.82 ± 0.0847 |
| | $V_{0.5,2}$ (mV) | -123 ± 3.10 | -120 ± 3.09 | -123 ± 1.76 | -123 ± 2.74 | -121 ± 1.80 | -122 ± 1.19 | -123 ± 0.923 | -125 ± 1.65 |
| | $z_2$ ($e_0$) | 2.37 ± 0.177 | 2.53 ± 0.138 | 2.36 ± 0.103 | 2.11 ± 0.0987 | 2.12 ± 0.124 | 2.10 ± 0.124 | 1.90 ± 0.0979 | 1.81 ± 0.0386 |
| | $n$ (cells) | 8 | 10 | 24 | 12 | 11 | 11 | 9 | 17 |
| VSD-I<br>$\beta_3$<br>(Fig.3i) | $V_h$ (mV) | -120 | -100 | -90 | -80 | -60 | -40 | 40 | |
| | $F_1$ (%) | 97.0 ± 1.70 | 73.4 ± 4.61 | 46.8 ± 6.76 | 33.1 ± 3.33 | 17.6 ± 3.41 | 1.06 ± 1.06 | 0.00 ± 0.00 | |
| | $V_{0.5,1}$ (mV) | -5.47 ± 0.920 | -5.72 ± 0.884 | -5.25 ± 1.44 | -5.27 ± 0.629 | -5.08 ± 0.865 | -5.00 ± 1.04 | - | |
| | $z_1$ ($e_0$) | 3.39 ± 0.147 | 3.44 ± 0.145 | 3.25 ± 0.173 | 4.08 ± 0.228 | 4.67 ± 0.289 | 5.04 ± 0.258 | - | |
| | $V_{0.5,2}$ (mV) | -126 ± 2.00 | -126 ± 1.84 | -125 ± 2.17 | -124 ± 1.35 | -124 ± 2.32 | -121 ± 0.783 | -121 ± 1.28 | |
| | $z_2$ ($e_0$) | 1.70 ± 0.110 | 1.70 ± 0.102 | 1.73 ± 0.110 | 1.75 ± 0.0760 | 1.57 ± 0.0881 | 1.60 ± 0.0873 | 1.76 ± 0.0675 | |
| | $n$ (cells) | 12 | 13 | 6 | 20 | 11 | 7 | 9 | |
| VSD-III<br>$\beta_{2a}$<br>(Fig.4a) | $V_h$ (mV) | -160 | -120 | -100 | -80 | -60 | -40 | 40 | |
| | $F_1$ (%) | 93.0 | 87.7 ± 2.14 | 77.7 ± 4.77 | 38.1 ± 3.08 | 25.9 ± 4.37 | 26.0 ± 4.05 | 0.233 ± 0.233 | |
| | $V_{0.5,1}$ (mV) | -13.1 | -16.2 ± 0.980 | -15.7 ± 1.33 | -15.9 ± 0.941 | -18.3 ± 1.15 | -12.5 ± 1.75 | -15.9 ± 0.941 | |
| | $z_1$ ($e_0$) | 2.87 | 3.33 ± 0.184 | 3.59 ± 0.236 | 3.28 ± 0.168 | 3.30 ± 0.26 | 3.95 ± 0.377 | 3.28 ± 0.168 | |
| | $V_{0.5,2}$ (mV) | -76.4 | -72.3 ± 1.14 | -71.5 ± 1.50 | -72.7 ± 1.14 | -73.1 ± 2.43 | -69.5 ± 1.34 | -72.7 ± 1.14 | |
| | $z_2$ ($e_0$) | 1.82 | 1.84 ± 0.131 | 2.09 ± 0.158 | 1.84 ± 0.120 | 1.95 ± 0.110 | 2.25 ± 0.328 | 1.84 ± 0.120 | |
| | $n$ (cells) | 2 | 17 | 11 | 19 | 6 | 5 | 19 | |
| VSD-III<br>$\beta_3$<br>(Fig.4d) | $V_h$ (mV) | -120 | -100 | -80 | -20 | 40 | | | |
| | $F_1$ (%) | 70.1 ± 2.62 | 33.4 ± 1.77 | 10.9 ± 1.90 | 0.00 ± 0.00 | 0.00 ± 0.00 | | | |
| | $V_{0.5,1}$ (mV) | -25.4 ± 1.56 | -26.0 ± 1.60 | -26.2 ± 1.64 | - | - | | | |
| | $z_1$ ($e_0$) | 2.77 ± 0.175 | 2.69 ± 0.167 | 2.70 ± 0.178 | - | - | | | |
| | $V_{0.5,2}$ (mV) | -77.7 ± 1.91 | -78.2 ± 2.03 | -78.9 ± 2.16 | -76.7 ± 1.72 | -78.3 ± 2.00 | | | |
| | $z_2$ ($e_0$) | 1.45 ± 0.0427 | 1.43 ± 0.0441 | 1.45 ± 0.0394 | 1.46 ± 0.0844 | 1.45 ± 0.0469 | | | |
| | $n$ (cells) | 12 | 11 | 13 | 6 | 11 | | | |
| VSD-IV<br>$\beta_{2a}$<br>(Fig.4g) | $V_h$ (mV) | -120 | -100 | -80 | -60 | -20 | 40 | | |
| | $F_1$ (%) | 55.4 ± 2.94 | 52.1 ± 5.39 | 37.6 ± 2.26 | 32.2 ± 1.59 | 2.80 ± 1.43 | 0.00 ± 0.00 | | |
| | $V_{0.5,1}$ (mV) | -9.58 ± 0.922 | -7.73 ± 0.917 | -8.59 ± 0.737 | -9.63 ± 1.22 | -7.96 ± 1.10 | - | | |
| | $z_1$ ($e_0$) | 2.97 ± 0.183 | 3.57 ± 0.324 | 3.17 ± 0.189 | 2.48 ± 0.0905 | 3.68 ± 0.223 | - | | |
| | $V_{0.5,2}$ (mV) | -99.0 ± 2.65 | -93.0 ± 2.65 | -97.2 ± 1.95 | -102 ± 2.36 | -93.9 ± 1.63 | -99.2 ± 2.38 | | |
| | $z_2$ ($e_0$) | 0.656 ± 0.0318 | 0.617 ± 0.0783 | 0.665 ± 0.0267 | 0.641 ± 0.0414 | 0.679 ± 0.0597 | 0.695 ± 0.0292 | | |
| | $n$ (cells) | 18 | 4 | 26 | 4 | 7 | 19 | | |
| VSD-IV<br>$\beta_3$<br>(Fig.4j) | $V_h$ (mV) | -160 | -120 | -100 | -90 | -80 | -60 | -40 | 40 |
| | $F_1$ (%) | 69.2 ± 1.52 | 62.4 ± 2.48 | 50.6 ± 4.07 | 39.1 ± 3.36 | 27.6 ± 2.04 | 12.8 ± 3.27 | 4.02 ± 2.28 | 0.00 ± 0.00 |
| | $V_{0.5,1}$ (mV) | -14.1 ± 0.369 | -15.3 ± 0.701 | -16.1 ± 0.822 | -14.7 ± 0.682 | -15.0 ± 0.533 | -16.1 ± 0.822 | -14.4 ± 1.14 | - |
| | $z_1$ ($e_0$) | 2.91 ± 0.275 | 2.98 ± 0.191 | 3.24 ± 0.251 | 2.64 ± 0.191 | 2.97 ± 0.153 | 3.24 ± 0.251 | 2.68 ± 0.253 | - |
| | $V_{0.5,2}$ (mV) | -102 ± 2.18 | -100 ± 0.772 | -101 ± 1.23 | -101 ± 1.20 | -101 ± 0.785 | -101 ± 1.23 | -99.5 ± 0.714 | -101 ± 0.785 |
| | $z_2$ ($e_0$) | 0.787 ± 0.0193 | 0.773 ± 0.0181 | 0.763 ± 0.0261 | 0.780 ± 0.0204 | 0.776 ± 0.0140 | 0.763 ± 0.0261 | 0.784 ± 0.0271 | 0.776 ± 0.0140 |
| | $n$ (cells) | 4 | 11 | 6 | 7 | 15 | 6 | 5 | 15 |

**Table. S2.**
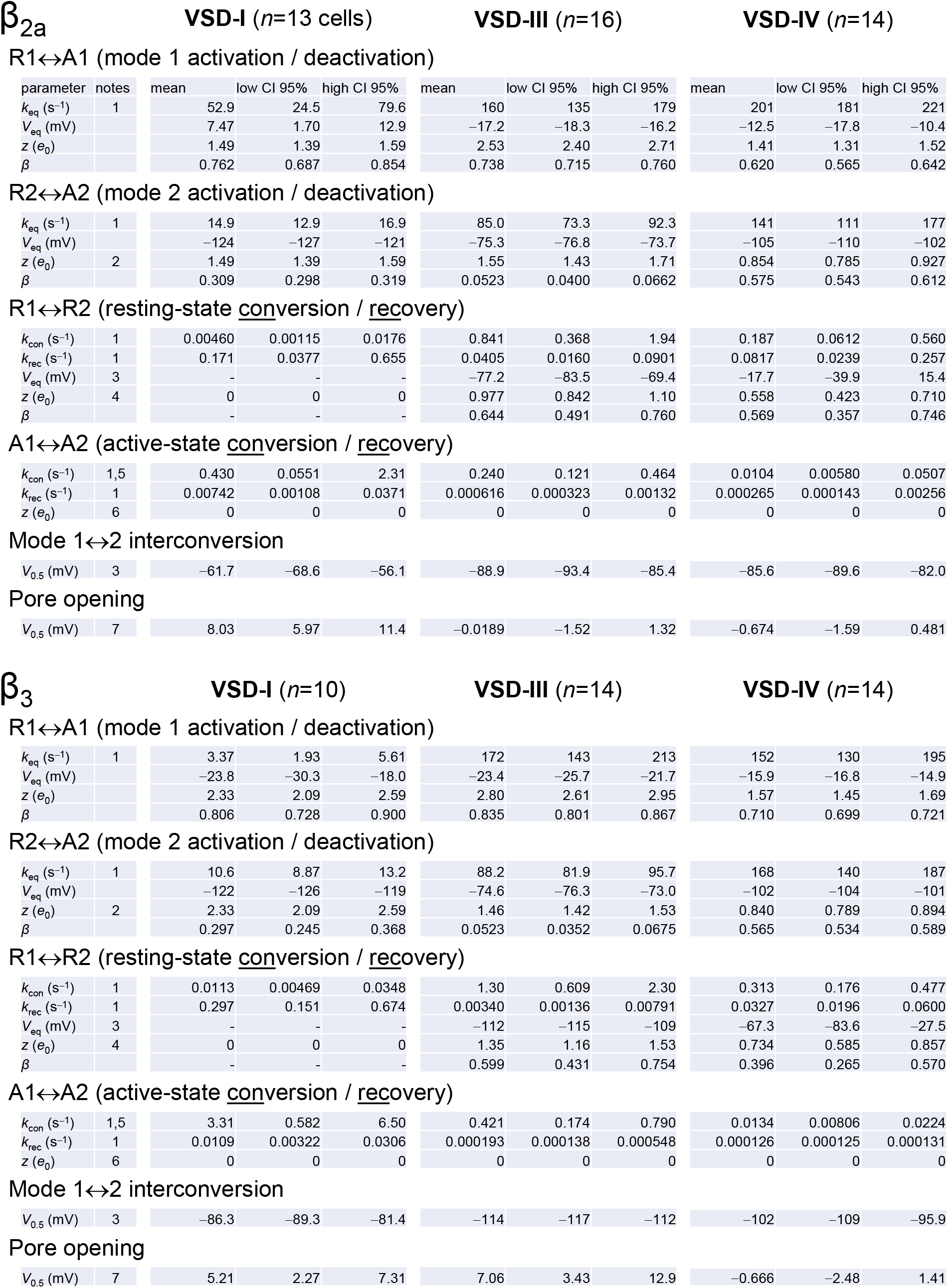
4-state model kinetic parameters. Notes: 1: geometric mean; 2: constrained parameter (charge conservation; eq.10); 3: calculated after fitting; 4: fixed parameter (VSD-I only); 5: constrained parameter (microscopic reversibility; eq.9); 6: fixed parameter (all VSDs); 7: from Boltzmann fits of tail currents (eq.1) at *V*_h_ = −80 mV.

## References

1. Mori Y, et al. Primary structure and functional expression from complementary DNA of a brain calcium channel. Nature 350, 398–402 (1991).

2. Inchauspe CG, Martini FJ, Forsythe ID, Uchitel OD. Functional compensation of P/Q by N-type channels blocks short-term plasticity at the calyx of Held presynaptic terminal. J Neurosci 24, 10379–10383 (2004).

3. Li L, Bischofberger J, Jonas P. Differential gating and recruitment of P/Q-, N-, and R-type Ca2+ channels in hippocampal mossy fiber boutons. J Neurosci 27, 13420–13429 (2007).

4. Kulik Á, et al. Immunocytochemical localization of the α1A subunit of the P/Q-type calcium channel in the rat cerebellum. European Journal of Neuroscience 19, 2169–2178 (2004).

5. Westenbroek R, et al. Immunochemical identification and subcellular distribution of the alpha 1A subunits of brain calcium channels. The Journal of Neuroscience 15, 6403–6418 (1995).

6. Zamponi GW, Striessnig J, Koschak A, Dolphin AC. The Physiology, Pathology, and Pharmacology of Voltage-Gated Calcium Channels and Their Future Therapeutic Potential. Pharmacol Rev 67, 821–870 (2015).

7. Dolphin AC, Lee A. Presynaptic calcium channels: specialized control of synaptic neurotransmitter release. Nat Rev Neurosci 21, 213–229 (2020).

8. Klein M, Shapiro E, Kandel ER. Synaptic plasticity and the modulation of the Ca2+ current. J Exp Biol 89, 117–157 (1980).

9. Tsodyks MV, Markram H. The neural code between neocortical pyramidal neurons depends on neurotransmitter release probability. Proc Natl Acad Sci U S A 94, 719–723 (1997).

10. Patil PG, Brody DL, Yue DT. Preferential closed-state inactivation of neuronal calcium channels. Neuron 20, 1027–1038 (1998).

11. Forsythe ID, Tsujimoto T, Barnes-Davies M, Cuttle MF, Takahashi T. Inactivation of presynaptic calcium current contributes to synaptic depression at a fast central synapse. Neuron 20, 797–807 (1998).

12. Liu Z, Ren J, Murphy TH. Decoding of synaptic voltage waveforms by specific classes of recombinant high-threshold Ca2+ channels. The Journal of Physiology 553, 473–488 (2003).

13. Pietrobon D. CaV2.1 channelopathies. Pflügers Archiv - European Journal of Physiology 460, 375–393 (2010).

14. Nanou E, Scheuer T, Catterall WA. Calcium sensor regulation of the Ca<sub>V</sub>2.1 Ca<sup>2+</sup> channel contributes to long-term potentiation and spatial learning. Proceedings of the National Academy of Sciences 113, 13209–13214 (2016).

15. Nanou E, Catterall WA. Calcium Channels, Synaptic Plasticity, and Neuropsychiatric Disease. Neuron 98, 466–481 (2018).

16. Borjesson SI, Elinder F. Structure, function, and modification of the voltage sensor in voltage-gated ion channels. Cell Biochem Biophys 52, 149–174 (2008).

17. Pantazis A, Savalli N, Sigg D, Neely A, Olcese R. Functional heterogeneity of the four voltage sensors of a human L-type calcium channel. Proc Natl Acad Sci U S A 111, 18381–18386 (2014).

18. Taglialatela M, Toro L, Stefani E. Novel voltage clamp to record small, fast currents from ion channels expressed in Xenopus oocytes. Biophys J 61, 78–82 (1992).

19. Stefani E, Bezanilla F. Cut-open oocyte voltage-clamp technique. Methods Enzymol 293, 300–318 (1998).

20. Pantazis A, Olcese R. Cut-Open Oocyte Voltage-Clamp Technique. In: Encyclopedia of Biophysics (eds Roberts G, Watts A). Springer (2019).

21. Mannuzzu LM, Moronne MM, Isacoff EY. Direct physical measure of conformational rearrangement underlying potassium channel gating. Science 271, 213–216 (1996).

22. Gandhi CS, Olcese R. The voltage-clamp fluorometry technique. Methods Mol Biol 491, 213–231 (2008).

23. Priest M, Bezanilla F. Functional Site-Directed Fluorometry. Adv Exp Med Biol 869, 55–76 (2015).

24. Bhat S, Blunck R. Characterising ion channel structure and dynamics using fluorescence spectroscopy techniques. Biochem Soc Trans 50, 1427–1445 (2022).

25. Stea A, et al. Localization and functional properties of a rat brain alpha 1A calcium channel reflect similarities to neuronal Q- and P-type channels. Proc Natl Acad Sci U S A 91, 10576–10580 (1994).

26. De Waard M, Campbell KP. Subunit regulation of the neuronal alpha 1A Ca2+ channel expressed in Xenopus oocytes. J Physiol 485 (Pt 3), 619–634 (1995).

27. Savalli N, Kondratiev A, Toro L, Olcese R. Voltage-dependent conformational changes in human Ca(2+)- and voltage-activated K(+) channel, revealed by voltage-clamp fluorometry. Proc Natl Acad Sci U S A 103, 12619–12624 (2006).

28. Pantazis A, Kohanteb AP, Olcese R. Relative motion of transmembrane segments S0 and S4 during voltage sensor activation in the human BK(Ca) channel. J Gen Physiol 136, 645–657 (2010).

29. Pantazis A, Olcese R. Relative transmembrane segment rearrangements during BK channel activation resolved by structurally assigned fluorophore-quencher pairing. J Gen Physiol 140, 207–218 (2012).

30. Pantazis A, Westerberg K, Althoff T, Abramson J, Olcese R. Harnessing photoinduced electron transfer to optically determine protein sub-nanoscale atomic distances. Nat Commun 9, 4738 (2018).

31. Li Z, et al. Structural basis for different ω-agatoxin IVA sensitivities of the P-type and Q-type Ca(v)2.1 channels. Cell Res 34, 455–457 (2024).

32. Brum G, Rios E. Intramembrane charge movement in frog skeletal muscle fibres. Properties of charge 2. J Physiol 387, 489–517 (1987).

33. Shirokov R, Levis R, Shirokova N, Rios E. Two classes of gating current from L-type Ca channels in guinea pig ventricular myocytes. J Gen Physiol 99, 863–895 (1992).

34. Olcese R, Latorre R, Toro L, Bezanilla F, Stefani E. Correlation between charge movement and ionic current during slow inactivation in Shaker K+ channels. J Gen Physiol 110, 579–589 (1997).

35. Mannikko R, Pandey S, Larsson HP, Elinder F. Hysteresis in the voltage dependence of HCN channels: conversion between two modes affects pacemaker properties. J Gen Physiol 125, 305–326 (2005).

36. Stotz SC, Jarvis SE, Zamponi GW. Functional roles of cytoplasmic loops and pore lining transmembrane helices in the voltage-dependent inactivation of HVA calcium channels. J Physiol 554, 263–273 (2004).

37. Dong Y, et al. Closed-state inactivation and pore-blocker modulation mechanisms of human Ca(V)2.2. Cell Rep 37, 109931 (2021).

38. Gao Y, et al. Molecular insights into the gating mechanisms of voltage-gated calcium channel Ca(V)2.3. Nat Commun 14, 516 (2023).

39. Yao X, et al. Structures of the R-type human Ca(v)2.3 channel reveal conformational crosstalk of the intracellular segments. Nat Commun 13, 7358 (2022).

40. Miller JA, Constantinidis C. Timescales of learning in prefrontal cortex. Nat Rev Neurosci 25, 597–610 (2024).

41. Nilsson M, et al. Voltage-dependent G-protein regulation of Ca(V)2.2 (N-type) channels. Sci Adv 10, eadp6665 (2024).

42. Gao S, Yao X, Yan N. Structure of human Ca(v)2.2 channel blocked by the painkiller ziconotide. Nature 596, 143–147 (2021).

43. Savalli N, Pantazis A, Sigg D, Weiss JN, Neely A, Olcese R. The alpha2delta-1 subunit remodels CaV1.2 voltage sensors and allows Ca2+ influx at physiological membrane potentials. J Gen Physiol 148, 147–159 (2016).

44. Savalli N, et al. The distinct role of the four voltage sensors of the skeletal CaV1.1 channel in voltage-dependent activation. J Gen Physiol 153, (2021).

45. Van Petegem F, Clark KA, Chatelain FC, Minor DL, Jr. Structure of a complex between a voltage-gated calcium channel beta-subunit and an alpha-subunit domain. Nature 429, 671–675 (2004).

46. Colecraft HM, Patil PG, Yue DT. Differential occurrence of reluctant openings in G-protein-inhibited N- and P/Q-type calcium channels. J Gen Physiol 115, 175–192 (2000).

47. Zamponi GW, Currie KP. Regulation of Ca(V)2 calcium channels by G protein coupled receptors. Biochim Biophys Acta 1828, 1629–1643 (2013).

48. Lee SR, Adams PJ, Yue DT. Large Ca(2)(+)-dependent facilitation of Ca(V)2.1 channels revealed by Ca(2)(+) photo-uncaging. J Physiol 593, 2753–2778 (2015).

49. Lee A, Westenbroek RE, Haeseleer F, Palczewski K, Scheuer T, Catterall WA. Differential modulation of Ca(v)2.1 channels by calmodulin and Ca2+-binding protein 1. Nat Neurosci 5, 210–217 (2002).

50. Perni S, Beam K. Neuronal junctophilins recruit specific Ca(V) and RyR isoforms to ER-PM junctions and functionally alter Ca(V)2.1 and Ca(V)2.2. Elife 10, (2021).

51. Bezprozvanny I, Scheller RH, Tsien RW. Functional impact of syntaxin on gating of N-type and Q-type calcium channels. Nature 378, 623–626 (1995).

52. Shih TM, Smith RD, Toro L, Goldin AL. High-level expression and detection of ion channels in Xenopus oocytes. Methods Enzymol 293, 529–556 (1998).

53. Colquhoun D, Hawkes AG. Relaxation and fluctuations of membrane currents that flow through drug-operated channels. Proc R Soc Lond B Biol Sci 199, 231–262 (1977).

54. Colquhoun D, Hawkes AG. On the stochastic properties of bursts of single ion channel openings and of clusters of bursts. Philos Trans R Soc Lond B Biol Sci 300, 1–59 (1982).

55. Acerbi L, Ma WJ. Practical Bayesian Optimization for Model Fitting with Bayesian Adaptive Direct Search. Adv Neur In 30, (2017).

56. Goddard TD, et al. UCSF ChimeraX: Meeting modern challenges in visualization and analysis. Protein Sci 27, 14–25 (2018).

57. Pettersen EF, et al. UCSF ChimeraX: Structure visualization for researchers, educators, and developers. Protein Sci 30, 70–82 (2021).

58. Meng EC, et al. UCSF ChimeraX: Tools for structure building and analysis. Protein Sci 32, e4792 (2023).

